# Aminoacyl-tRNA synthetase evolution within the dynamic tripartite translation system of plant cells

**DOI:** 10.1101/2022.12.12.520092

**Authors:** Daniel B. Sloan, Rachael A. DeTar, Jessica M. Warren

## Abstract

Eukaryotes maintain separate protein translation systems for nuclear and organellar genes, including distinct sets of tRNAs and aminoacyl-tRNA synthetases (aaRSs). In animals, mitochondrial-targeted aaRSs are expressed at lower levels and are less conserved in sequence than cytosolic aaRSs involved in translation of nuclear mRNAs, likely reflecting lower translational demands in mitochondria. In plants, translation is further complicated by the presence of plastids, which share most aaRSs with mitochondria. In addition, plant mitochondrial tRNA pools have a dynamic history of gene loss and functional replacement by tRNAs from other compartments. To investigate the consequences of these distinctive features of translation in plants, we analyzed sequence evolution in angiosperm aaRSs. In contrast to previously studied eukaryotic systems, we found that plant organellar and cytosolic aaRSs exhibit only a small difference in expression levels, and organellar aaRSs are slightly *more* conserved than cytosolic aaRSs. We hypothesize that these patterns result from high translational demands associated with photosynthesis in mature chloroplasts. We also investigated aaRS evolution in *Sileneae*, an angiosperm lineage with extensive mitochondrial tRNA replacement and aaRS retargeting. We predicted positive selection for changes in aaRS sequence resulting from these recent changes in subcellular localization and tRNA substrates but found little evidence for accelerated sequence divergence. Overall, the complex tripartite translation system in plant cells appears to have imposed more constraints on the long-term evolutionary rates of organellar aaRSs compared to other eukaryotic lineages, and plant aaRS protein sequences appear largely robust to more recent perturbations in subcellular localization and tRNA interactions.

## INTRODUCTION

Eukaryotic cells contain multiple genomic compartments, reflecting their endosymbiotic origins. Accordingly, separate translation systems exist for production of proteins encoded in nuclear, mitochondrial (mt), and plastid (pt) genomes. Organellar translation systems retain bacterial-like ribosomes (Harris, et al. 1994; Smits, et al. 2007) and often rely on ancestral mt-tRNA genes that still reside in the mt genome (Salinas-Giegé, et al. 2015). However, much of the organellar translation machinery is now encoded in the nuclear genome (Timmis, et al. 2004; Giannakis, et al. 2022). In most eukaryotes, this includes all of the aminoacyl-tRNA synthetases (aaRSs), which are responsible for recognizing tRNAs and charging them with the correct amino acid. Most eukaryotes express two largely distinct sets of aaRSs – one that is responsible for translation of nuclear mRNAs in the cytosol and a second organellar set that functions in mitochondria (and plastids in lineages such as plants) (Brindefalk, et al. 2007; Duchêne, et al. 2009; Salinas-Giegé, et al. 2015). The direct functional interactions between gene products encoded in different genomes within eukaryotic cells can lead to cytonuclear coadaptation (Rand, et al. 2004; Hill 2015; Sloan, et al. 2018). The rapid sequence divergence observed in many mt genomes may select for coevolutionary responses in the nucleus (Osada and Akashi 2012; Havird, et al. 2015; Barreto, et al. 2018), and incompatibilities between mt tRNAs and nuclear-encoded aaRSs can have severe fitness consequences (Meiklejohn, et al. 2013).

In animals, mt aaRSs exhibit faster sequence divergence than their cytosolic counterparts (Pett and Lavrov 2015; Adrion, et al. 2016; Barreto, et al. 2018). One possible explanation is that selection on nuclear-encoded aaRSs to respond to changes in mt tRNA genes has caused these higher evolutionary rates. However, mt aaRSs also have lower expression levels than cytosolic aaRSs, which may indicate that they are simply under weaker purifying selection because translational demands are less intense in mitochondria than in the cytosol (Sloan, et al. 2014; Pett and Lavrov 2015). Indeed, when accounting for gene expression levels, there is often no detectable difference in evolutionary rates between mt and cytosolic aaRSs in animals (Adrion, et al. 2016; Barreto, et al. 2018). Therefore, the evidence that mitonuclear coevolution is a substantial contributor to rates of mt aaRS sequence evolution remains very limited.

In contrast to the detailed work in animal systems, evolutionary rates for organellar and cytosolic aaRSs have not been studied extensively in plants. Yet, there are reasons to expect that the presence of plastids may place distinct constraints on plants organellar aaRS evolution. Pt translation is responsible for massive levels of protein production. The chloroplasts themselves house ~80% of the total protein content in leaf mesophyll cells (Heinemann, et al. 2021). Although this total also includes the large number of nuclear-encoded proteins that are imported into chloroplasts from the cytosol, pt genes produce the majority of mRNA transcripts in the cell (Forsythe, et al. 2022), and their protein products such as PsbA and RbcL have exceptionally high rates of translation (Chotewutmontri and Barkan 2018). There are two largely separate classes of aaRSs in plants: cytosolic and organellar, with the latter being dual-targeted to mitochondria and plastids (Duchêne, et al. 2005; Duchêne, et al. 2009). Therefore, translational demands in the plastid are expected to affect conservation and rates of evolution of aaRSs with shared function in the mitochondria.

The complex and dynamic mixture of tRNAs that function in plant mitochondria adds another potential complexity to the evolution of plant aaRSs. Plant mt genomes contain tRNA genes from multiple origins, including a portion of the ancestral mt gene set as well as horizontal transfers from the pt genome, other bacterial genomes, and the mt genomes of other plants (Small, et al. 1999; Warren and Sloan 2020). However, even this heterogeneous set of tRNA genes is insufficient for translation of all codons, and plant mitochondria import additional nuclear-encoded tRNAs from the cytosol (Michaud, et al. 2011). The extent of cytosolic import varies among plant species as there is recent and ongoing mt tRNA gene loss in many lineages. The angiosperm tribe *Sileneae* is a striking example, with some species retaining as many as 14 mt tRNA genes while other close relatives have only two or three (Warren, et al. 2021).

The history of plant mt tRNA gene loss raises questions about effects on aaRS function. Eukaryotic/nuclear tRNAs are essentially unrecognizable in primary sequence when compared to bacterial-like mt tRNAs. Therefore, it is not clear if and how organellar aaRS are able to charge these newly imported tRNAs in the mitochondria. Results from *in silico* analysis of putative targeting peptides and fluorescence microscopy assays of subcellular localization in *Silenenae* have identified two alternative pathways (Warren, et al. 2022). In some cases, changes in tRNA import have been accompanied by retargeting of the corresponding cytosolic aaRS such that the ancestral pairing between tRNA and aaRS is maintained and simply relocated to a second compartment. In these cases, there is also the potential for concomitant loss of mt targeting in the organellar aaRSs such that ancestral dual-targeted enzymes now function exclusively in plastids. In other cases of mt tRNA gene loss and functional replacement, there is no evidence of aaRS retargeting, implying that the ancestral organellar aaRS has evolved to charge a newly imported cytosolic tRNA. These two alternative responses to mt tRNA gene loss and functional replacement are also observed in more ancient examples in plant evolution (Duchêne, et al. 2009).

Such changes in targeting or tRNA substrates represent potentially radical perturbations in aaRS function, and we hypothesize that they could result in positive selection and accelerated evolution in aaRS sequence for at least three different reasons. First, aaRSs evolving to charge new tRNA substrates may require changes to key regions involved in recognition of tRNA “identity elements” (Giegé, et al. 1998; Igloi 2021, 2022). Second, mitochondria differ from the cytosol in numerous respects (e.g., osmotic conditions) that could potentially affect protein folding and tRNA interactions and, thus, create selection for changes in protein sequence. Third, organellar aaRSs that ancestrally functioned in both the mitochondria and plastids but lose mt targeting may experience altered selection pressures in specializing exclusively on plastid function. All these mechanisms would be predicted to accelerate evolution and produce signatures of positive selection for amino acid substitutions in lineages that have experienced recent changes in mt tRNA gene content.

In this study, we test predictions about how the distinctive features of plant organellar translation have shaped rates of aaRS evolution. We take a phylogenetic approach to analyze sequence evolution in plant aaRS families, including long-term patterns across divergent angiosperms and the more recent history of divergence among *Sileneae* species that differ greatly in mt-tRNA gene content and cytosolic import.

## RESULTS

### Angiosperm organellar aaRSs exhibit a high degree of sequence conservation

To characterize long-term rates of aaRS amino-acid sequence divergence in angiosperms, we generated trees for each aaRS with representatives of eudicots (*Arabidopsis thaliana* and *Vitis vinifera*) and monocots (*Oryza sativa* and *Spirodela polyrhiza*). Total sequence divergence (substitutions per site) was determined based on summed branch lengths within the tree (using an averaging approach in case of duplicated gene copies). We found that angiosperm organellar aaRS sequences evolved slightly (but significantly) slower than their cytosolic counterparts (*p* = 0.0178; paired *t-*test; Figure 1). Although the magnitude of the difference between these two groups was small (25% lower rates for organellar aaRSs on average), this result presents a striking contrast with previous work in animals, which has shown that mt aaRSs evolve much more rapidly than cytosolic aaRSs (Pett and Lavrov 2015; Adrion, et al. 2016; Barreto, et al. 2018).

**Figure 1.**
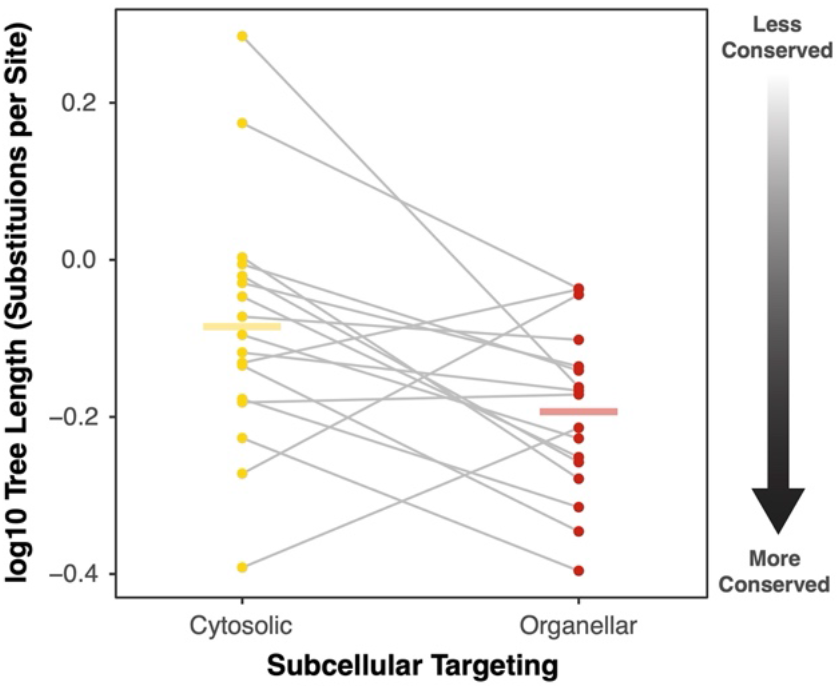
Lower rates of amino-acid substitutions in angiosperm organellar aaRSs than cytosolic aaRSs. Sequence divergence was quantified by summing branch lengths (measured as amino acid substitutions per site) from phylogenetic trees for each aaRS, containing orthologs from *A. thaliana*, *V. vinifera*, *O. sativa*, and *S. polyrhiza*. Points represent log10-transformed values for total tree length for each cytosolic (yellow) and organellar (red) aaRS, and gray lines connect pairs connect counterparts associated with the same amino acid. Horizontal bars represent the mean for each group. Organellar rates are significantly lower than cytosolic rates based on a paired *t-*test on log-transformed values (*p* = 0.0178).

### Angiosperm organellar aaRSs are expressed at only slightly lower levels than their cytosolic counterparts

One hypothesis to explain why mitochondrial aaRS sequences evolve rapidly in animals is that they are expressed at the lower levels than cytosolic aaRSs (Adrion, et al. 2016; Barreto, et al. 2018), and expression tends to be negatively correlated with rates of protein evolution as a general principle – the so-called E-R anticorrelation (Zhang and Yang 2015). Indeed, mRNA transcript abundance for mt aaRS has been shown to be approximately five-fold lower than for cytosolic aaRSs in multiple animal systems (Adrion, et al. 2016). Thus, we reasoned that plants may not show the same imbalance in expression level that is observed in animals. Using RNA-seq data from the EMBL-EBI Expression Atlas (Moreno, et al. 2022) from multiple *A. thaliana* tissue types, we found that plants exhibited lower expression levels for their organellar aaRS genes relative to their cytosolic counterparts (Figure 2). This difference is in the same direction as observed in animals. Therefore, expression level alone cannot fully explain the inverted relationship for aaRS substitution rates in organellar and cytosolic aaRSs for plants vs. animals. However, the differences in expression levels that we observed in *A. thaliana* (2.2-fold in flowers, 2.0-fold in leaves, and 1.5-fold in seedlings) are substantially smaller than those previously found in animal systems (Adrion, et al. 2016). This contrast suggests that plants maintain greater demands on organellar aaRS function than animals, which may contribute to their high degree of sequence conservation (Figure 2).

**Figure 2.**
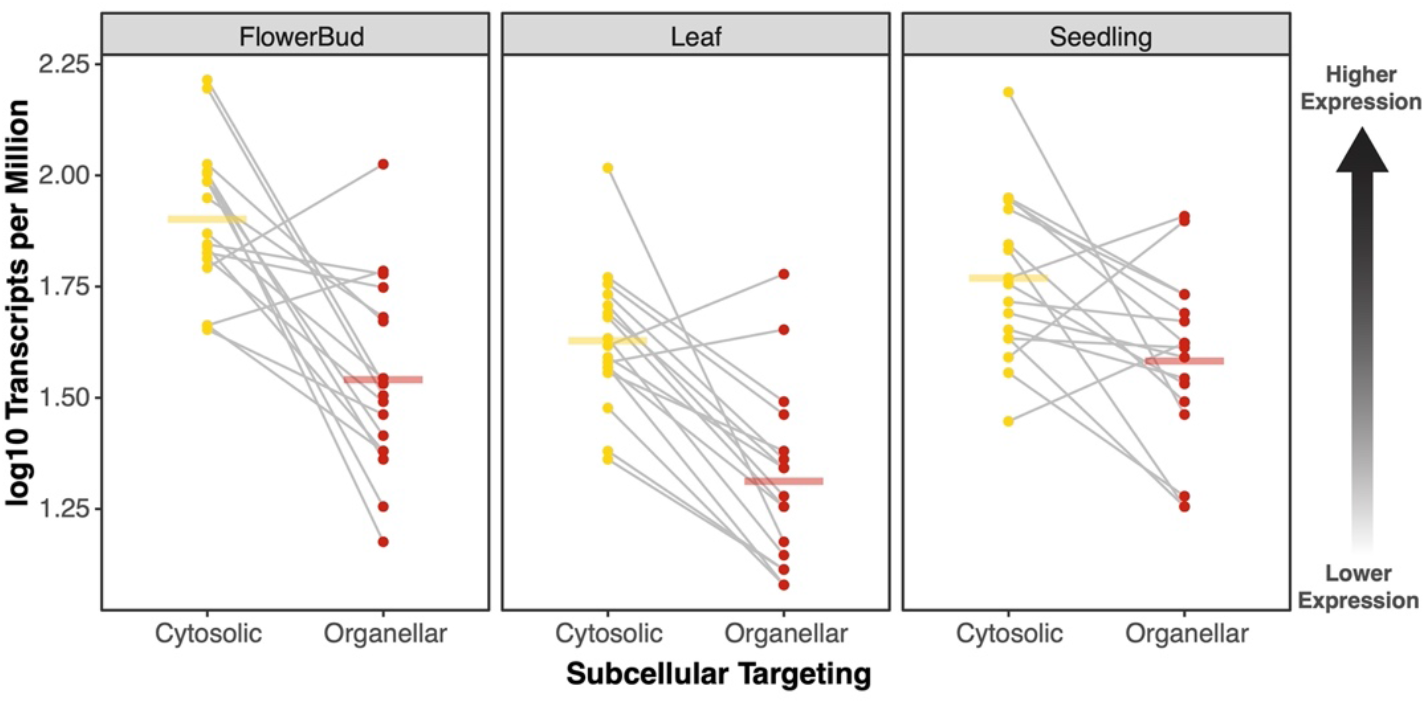
Lower levels of expression (mRNA transcript abundance) for organellar aaRS genes than cytosolic aaRS genes in three different *A. thaliana* tissue types based on data from the EMBL-EBI Expression Atlas. Points represent cytosolic (yellow) and organellar (red) aaRSs, and gray lines connect pairs connect counterparts associated with the same amino acid. Horizontal bars represent the mean for each group. A three-way ANOVA found significant effects of aaRS targeting (*p* = 1.2e-13), tissue (*p* = 1.5e-8), and aaRS amino-acid (*p* = 4.6e-4) on expression level as measured by log-transformed transcripts per million (TPM).

### Angiosperm organellar aaRSs are present in lower gene copy numbers than their cytosolic counterparts

Gene duplication is pervasive in plants (Panchy, et al. 2016), and the presence of duplicates potentially alters the selection pressures that can affect levels of sequence conservation (Lynch and Conery 2000). Therefore, we considered the possibility that organellar and cytosolic aaRSs gene families systematically differ in copy number. Using a previously generated phylogenomic sampling of 20 angiosperm species (Forsythe, et al. 2021), we compared the size of aaRS gene families and found that cytosolic families were 31% larger on average (*p* = 0.0006; paired *t*-test; Figure 3). Very similar results were obtained when the analysis was restricted to the 13 species for which whole genomes (and not just transcriptome assemblies) were available, with cytosolic families being 30% larger than organellar families (*p* = 0.0026; paired *t*-test). Therefore, the rate of gene duplication and/or retention appears to be higher for cytosolic aaRSs, potentially relaxing selection pressures on these gene copies. The larger number of copies for cytosolic aaRSs also parallels the very high level of paralogy for cytosolic ribosomal proteins relative to mt and pt ribosomal proteins (Yamaguchi and Subramanian 2000; Yamaguchi, et al. 2000; Barakat, et al. 2001; Bonen and Calixte 2006).

**Figure 3.**
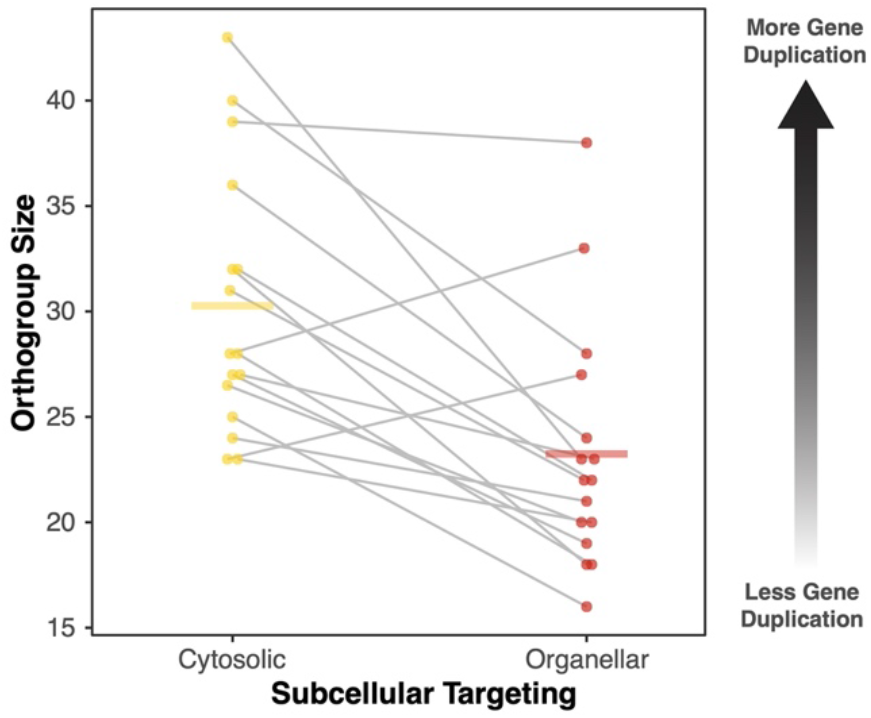
Higher rates of gene duplication/retention for cytosolic aaRSs than organellar aaRSs in angiosperms. Gene family size was assessed based on the number of genes in orthogroups from a previous analysis of angiosperm phylogenomic diversity (Forsythe, et al. 2021). Points represent cytosolic (yellow) and organellar (red) aaRSs, and gray lines connect pairs connect counterparts associated with the same amino acid. Horizontal bars represent the mean for each group. Cytosolic aaRS orthogroup size was significantly larger than organellar orthogroup size (*p* = 0.0006; paired *t*-test).

### Rates of aaRS sequence evolution in *Sileneae*

To assess whether recent changes in subcellular targeting or tRNA substrates for aaRSs in *Sileneae* had altered selection pressures and rates of sequence evolution, we partitioned aaRS gene trees to assign “foreground” and “background” branches. We defined foreground branches as those with an aaRS in one of the following categories: 1) an ancestral organellar aaRS inferred to charge a novel cytosolic tRNA substrate that is now imported into the mitochondria, 2) an ancestral cytosolic aaRS inferred to have gained mt targeting, or 3) an ancestral dual-targeted organellar aaRS inferred to have lost mt targeting and specialized on pt function.

We first tested the hypothesis that these foreground branches had experienced positive selection and accelerated evolution in the form of an increased ratio of non-synonymous to synonymous substitutions (*d*_N_/*d*_S_) by performing branch tests on each tree. These tests generally failed to detect evidence that foreground branches had experienced faster protein sequence evolution (Table 1). In two cases (organellar AspRS and cytosolic TyrRS), the nominal *p*-value for increased rates on foreground branches was less than 0.05, but these comparisons did not remain significant after correction for multiple tests.

**Table 1.** Branch tests for increased *d*_N_/*d*_S_ ratios in response to inferred changes in subcellular localization or tRNA substrate. Foreground branches for each test were assigned as described in the methods section.

| aaRS | Ancestral Targeting | Hypothesized Mechanism | Foreground $d_N/d_S$ | Background $d_N/d_S$ | p-value (raw) |
| --- | --- | --- | --- | --- | --- |
| AsnRS | Organellar | New Substrate | 0.14 | 0.08 | 0.1153 |
| AspRS | Cytosolic | Mito Import | 0.16 | 0.26 | 0.4166 |
| AspRS | Organellar | New Substrate | 0.28 | 0.17 | 0.0253 |
| CysRS | Organellar | New Substrate | 0.15 | 0.15 | 0.8875 |
| GlnRS | Cytosolic | Mito Import | 0.14 | 0.11 | 0.2301 |
| GluRS | Organellar | New Substrate | 0.31 | 0.16 | 0.0853 |
| HisRS | Organellar | New Substrate | 0.12 | 0.13 | 0.7518 |
| LysRS | Cytosolic | Mito Import | 0.13 | 0.09 | 0.1473 |
| MetRS | Organellar | Plastid Specialization | 0.27 | 0.26 | 1.0000 |
| PheRS | Organellar | New Substrate* | 0.14 | 0.12 | 0.6390 |
| ProRS | Cytosolic | Mito Import | 0.12 | 0.08 | 0.0719 |
| TrpRS | Organellar | Plastid Specialization | 0.27 | 0.14 | 0.0979 |
| TrpRS | Cytosolic | Mito Import | 0.10 | 0.17 | 0.1492 |
| TyrRS | Cytosolic | Mito Import | 0.34 | 0.23 | 0.0088 |

\*Organellar PheRS could also be a potential case of plastid specialization because *Sileneae* contains duplicate copies with one apparently functioning in the mitochondria and another apparently functioning in the plastids (Warren, et al. 2022), but this analysis was run with the putative mt copy set as the foreground branches.

Because branch tests average *d*_N_/*d*_S_ ratios across the entire length of a gene and may dilute the effects of positive selection acting on a small subset of codons within a gene, we also applied a suite of branch-site tests, which aim to detect positive selection even if it has favored amino acid substitutions at only a small number of sites: aBSREL (Smith, et al. 2015), BUSTED (Murrell, et al. 2015), and MEME (Murrell, et al. 2012). However, these tests also yielded little evidence of positive selection or accelerated evolution on foreground branches. aBSREL tests each foreground branch for evidence that at least some proportion of codons have evolved under positive selection and only identified a single branch in a single tree (*Silene vulgaris* 49961 in the cytosolic TyrRS tree) as significant, but the raw *p*-value of 0.0297 did not remain significant after correction for multiple tests (Table 2). BUSTED performs a similar but more sensitive analysis, assessing whether there is evidence for positive selection at any site on any foreground branch. This analysis also identified the cytosolic TyrRS tree as only one with evidence of any positive selection (Table 2). Finally, we used MEME to identify specific codons that may have experienced episodic positive selection on foreground branches, using a permissive (raw) significance threshold of *p*< 0.10. This analysis identified a relatively small number of codons as candidates for positive selection, again finding the most evidence of positive selection in the cytosolic TyrRS tree (Table 2). Therefore, cytosolic TyrRS clearly emerged as the strongest candidate for positive selection, but overall evidence for accelerated aaRS evolution in response to changes in subcellular targeting or tRNA substrates was limited.

**Table 2.** Summary of branch-site for increased *d*_N_/*d*_S_ ratios in response to inferred changes in subcellular localization or tRNA substrate. The trees and hypothesized mechanisms are the same as described in Table 1. Foreground branches for each test were assigned as described in the methods section. All reported *p*-values are raw (i.e., uncorrected for multiple tests). Reported codon from MEME analyses are based on a raw significance threshold of *p*< 0.1.

| aaRS | Ancestral Targeting | aBSREL Branches (p-value) | BUSTED p-value | MEME Codon Position(s) |
| --- | --- | --- | --- | --- |
| AsnRS | Organellar | . | 0.4165 | . |
| AspRS | Cytosolic | . | 0.0997 | 88;89 |
| AspRS | Organellar | . | 0.3811 | 7;141;370;439;582 |
| CysRS | Organellar | . | 0.5000 | 15;424 |
| GlnRS | Cytosolic | . | 0.1150 | 226;752;776 |
| GluRS | Organellar | . | 0.5000 | 83;311;385 |
| HisRS | Organellar | . | 0.5000 | . |
| LysRS | Cytosolic | . | 0.4042 | 242;539;581 |
| MetRS | Organellar | . | 0.3507 | 181;516;543 |
| PheRS | Organellar | . | 0.3601 | 175;225;363;373 |
| ProRS | Cytosolic | . | 0.2282 | 346;433 |
| TrpRS | Organellar | . | 0.5000 | . |
| TrpRS | Cytosolic | . | 0.5000 | . |
| TyrRS | Cytosolic | *S. vulgaris* 49961 (0.0297) | 0.0006 | 28;57;95;138;197;361;479;667;731;738 |

## DISCUSSION

### The translational demands of photosynthesis and their effects on rates of organellar aaRS sequence evolution

An overall picture emerging from our analysis is that plant organellar aaRSs evolve under more intense constraints relative to cytosolic aaRSs than their counterparts in animals (Pett and Lavrov 2015; Adrion, et al. 2016; Barreto, et al. 2018). We hypothesize that this difference reflects the fact that plant organellar aaRSs must also function in plastids (Duchêne, et al. 2005) and that photosynthesis creates massive demands on plastid gene expression and translational systems (Chotewutmontri and Barkan 2018; Heinemann, et al. 2021; Forsythe, et al. 2022). Therefore, it is likely that organellar aaRSs are under unusually strong purifying selection in plants to function efficiently.

An alternative hypothesis is that plant aaRSs experience less positive selection than animal mt aaRSs. The mt genomes of bilaterian animals evolve much more rapidly than organellar genomes in plants (Wolfe, et al. 1987) and encode tRNAs that are unusually divergent in structure (Salinas-Giegé, et al. 2015). Thus, it is possible that rapid evolution in animal mt aaRSs results from selection for coevolutionary responses to changes in their cognate mt tRNAs, whereas such selections pressures would be largely absent for plant organellar aaRSs because of the slow sequence evolution that is typical of plant mt and pt tRNAs. However, support this hypothesis seems limited. If individual mutations in animal mt tRNAs are a driver of coevolutionary responses and rapid evolution in mt aaRSs, then functionally replacing a mt tRNA with an anciently divergent cytosolic counterpart would be expected to select for extensive changes in aaRS sequence. However, our analysis of the *Sileneae* organellar aaRS that appear to have adapted to charge newly imported cytosolic tRNAs found little or no evidence for accelerated aaRS evolution (Tables 1 and 2). Likewise, previous studies have cast doubt on the hypothesis that mitonuclear coevolution is primarily responsible for the rapid evolution of animal mt aaRSs; in particular, the low expression levels of mt aaRS genes may be a more important contributor to relaxed constraints on sequence evolution (Pett and Lavrov 2015; Adrion, et al. 2016; Barreto, et al. 2018). Our analysis of transcript abundance for plant aaRS genes provided partial support for this role of gene expression. Although, we found that plants did have lower expression levels for organellar aaRSs than for cytosolic aaRSs (Figure 2), the gap was substantially smaller than observed in animals, which may contribute to why plant organellar aaRSs do not evolve more slowly than their cytosolic counterparts (Figure 1).

Because most plant organellar aaRSs are dual-targeted (Duchêne, et al. 2005), it is difficult to decouple the effects of mt and pt function on constraining their sequence evolution. However, some clues may come from comparing other components of organellar translation machinery that are distinct between mitochondria and plastids. For example, unlike organellar aaRSs, there are separate sets of ribosomal proteins for mt and pt translation rather than a single set of dual-targeted proteins. In accordance with the hypothesis that pt translational demands impose more functional constraint than mt translation, *Arabidopsis* pt ribosomal protein genes exhibit rates of nonsynonymous sequence divergence that are much lower than those in mt ribosomal protein genes but statistically indistinguishable from those in cytosolic ribosomal protein genes (after excluding the rapidly evolving transit peptide sequences from mt- and pt-targeted proteins) (Sloan, et al. 2014).

One testable prediction of the hypothesis that translational demands associated with photosynthesis are the primary source of the selection constraining plant organellar aaRS sequence evolution is that heterotrophic plants that perform little or no photosynthesis (Wicke, et al. 2013) will show faster evolutionary rates for organellar aaRSs than for cytosolic aaRSs – more akin to observations in animals. We are currently investigating the evolution of aaRS sequence evolution and subcellular targeting in parasitic plants and their photoautotrophic relatives to test this prediction.

### Minimal effects of changes in subcellular targeting and tRNA substrates on the rate of aaRS sequence evolution

Despite the recent and major changes in subcellular targeting of tRNAs and aaRSs in *Sileneae* (Warren, et al. 2021; Warren, et al. 2022), we found little evidence that this rewiring of tRNA interaction networks has created positive selection for changes in aaRS protein sequence (Tables 1 and 2). One possible explanation for this apparently limited effect is that *Sileneae* organellar aaRS enzymes that have evolved to charge a newly imported cytosolic tRNA substrate were already preadapted to successfully recognize these tRNAs and, thus, required few changes to enzyme sequence. For example, we previously showed that mt and cytosolic tRNAs already shared key identity elements in most cases where the ancestral organellar aaRS apparently retained mt function upon import of a cytosolic tRNA (Warren, et al. 2022). We also found very limited evidence of positive selection in response to correlated retargeting of cytosolic tRNAs and aaRSs to the mitochondria. This may simply mean that the change in subcellular environment (cytosol vs. mitochondria) does not create strong selection for change in aaRS protein sequence because the ancestral aaRS-tRNA charging relationship is unchanged in these cases.

We should also note that some forms of positive selection may be difficult to detect with existing methods. Many of the current approaches to detect site-level positive selection were devised in the context of antagonistic coevolution, such as host-pathogen interactions, where selection for recurring amino-acid substitutions at the same position(s) might be expected. However, in other forms of positive selection, a single substitution may be sufficient to improve and stabilize molecular function (Hughes 2007). Therefore, future work could more directly test whether recent changes in *Sileneae* aaRS sequences have altered their tRNA specificity by using *in vitro* charging assays. Likewise, genome editing approaches could be used to assess whether the ancestral cytosolic aaRS enzyme bodies are interchangeable with those of the *Sileneae* aaRSs that have since been retargeted to the mitochondria. Such approaches may be able to detect more subtle finetuning that is required to maintain function in response to perturbations to subcellular localization or tRNA substrates.

## METHODS

### Angiosperm aaRS sequence curation, alignment, trimming, and phylogenetic rate analysis

Gene identifiers and subcellular targeting data for *A. thaliana* aaRS sequences were taken from Duchêne, et al. (2005) and Warren and Sloan (2020). Three types of aaRSs were excluded for the following reasons, which prevented a clean comparison between organellar and cytosolic aaRSs. 1) The only cytosolic AlaRS is also targeted to the organelles, 2) the only organellar ArgRS is also targeted the cytosol, and 3) there is no organellar GlnRS because plant mitochondria and plastids typically use a bacterial-like indirect charging pathway that involves GluRS (Duchêne, et al. 2005; Pujol, et al. 2008). Four aaRSs (GlyRS, LeuRS, ThrRS, ValRS) that function in the cytosol but also have mt (but not pt) localization were classified as cytosolic for the purposes of this analysis, and the LeuRS with pt (but not mt) localization was classified as organellar. Amino acid sequences for aaRSs from *A. thaliana* and three other distantly related angiosperms (*O. sativa*, *S. polyrhiza*, and *V. vinifera*) were taken from “orthogroups” produced with OrthoFinder (Emms and Kelly 2015) in a previous study (Forsythe, et al. 2021). The relevant orthogroups were selected based on the presence of the *A. thaliana* gene identifiers described above.

Sequences for each aaRS gene family were aligned with MAFFT v7.453 using the --auto option. The resulting alignments were manually curated in Geneious (Kearse, et al. 2012) to remove partial-length sequences and previously identified pseudogenes (Duchêne, et al. 2005) and to merge fragmented gene models. Sequences were then realigned with MAFFT. The N-terminus of alignments for organellar aaRSs was trimmed to eliminate predicted transit peptides because these are known to evolve rapidly and are cleaved from the rest of the functional enzyme during the mt and/or pt import process. Trimming was performed at the alignment position corresponding to the cleavage site predicted for *A. thaliana* by TargetP v2.0 (Armenteros, et al. 2019). After transit-peptide trimming, all alignments were manually inspected to trim poorly aligned regions at the N- and C-termini. In addition, poorly aligned internal regions in two sequences (*O. sativa* cytosolic TyrRS LOC_Os08g09260.1 and *V. vinifera* organellar PheRS GSVIVT01020339001) were also trimmed.

The resulting trimmed alignments were used for phylogenetic analysis with RAxML v8.2.12 (Stamatakis 2014) and PROTGAMMALG model of sequence evolution. To compare evolutionary rates among aaRSs, total tree length was calculated by summing individual branch lengths after averaging terminal branches for any paralogs to avoid inflating tree lengths for aaRSs with a history of gene duplication. Cytosolic PheRS is encoded as two separate subunits, so the total tree lengths for these subunits were averaged. Tree lengths for cytosolic vs. organellar aaRS were compared with a paired *t*-test on log-transformed values in R v4.0.5. Final trimmed alignments are available as supplemental material (File S1).

### aaRS gene copy number analysis

To test whether cytosolic and organellar aaRSs differed in gene copy number, we used the same orthogroup dataset described above (Forsythe, et al. 2021). The full dataset consists of 20 diverse angiosperm species, and we used the total number of sequences assigned to each orthogroup from all these species as a proxy for relative gene copy number. AlaRS, ArgRS, and GlnRS were excluded from this comparison for the same reasons as described above for the rate analysis. In addition, CysRS was excluded because the organellar and cytosolic targeting classes were assigned to the same orthogroup by OrthoFinder. Copy number values for the two cytosolic PheRS subunits were averaged. Copy numbers for cytosolic vs. organellar aaRS were compared with a paired *t*-test in R.

### *Arabidopsis thaliana* gene expression analysis

To analyze levels of aaRS gene expression (transcript abundance), data were obtained from the EMBL-EBI Expression Atlas (E-CURD-1: Araport 11 - RNA-seq of Arabidopsis thaliana Col-0 plants under different growth conditions from multiple studies) (Papatheodorou, et al. 2018). Three different tissue types were chosen from this dataset to be representative of diverse developmental stages:

- E-GEOD-30795: petal differentiation and expansion stage, long day length regimen, floral bud
- E-GEOD-44635: adult, long day length regimen, leaf
- E-MTAB-4242: seedling, long day length regimen, aerial part

Expression data were analyzed as transcripts per million (TPM). These TPM values were summed for paralogs. TPM values for the two cytosolic PheRS subunits were averaged. Log-transformed TPM data were analyzed with a 3-way ANOVA using the aov function in R, with targeting (cytosolic or organellar), tissue (flower bud, leaf, or seedling), and aaRS (AsnRS, AspRS, CysRS, etc.) as independent variables.

### *Sileneae* aaRS positive selection analysis

Alignments of trimmed *Sileneae* aaRS nucleotide sequences and phylogenetic trees were taken from a previous study (Warren, et al. 2022). The *A. thaliana* sequences that were originally used as outgroups were removed from the alignments. Coding sequences were then manually edited to be in frame and then aligned by codon (i.e., aligned with MAFFT as translated amino acid sequences and then back-translated to nucleotide sequences). Putative transit peptides for organellar aaRSs were trimmed based on predicted cleavage sites from TargetP.

To perform phylogenetic tests for selection on *Sileneae* aaRSs, *A. thaliana* genes were pruned from the original trees (Warren, et al. 2022), using the drop.tip function in the ape v5.4-1 package in R (Paradis and Schliep 2018). The phylotree.js tool (Shank, et al. 2018) was then used to label branches as “Foreground” if they were inferred to meet one of the criteria described in the main text or Table 1 (i.e., new tRNA substrate, gain of mt import by a cytosolic aaRS, or specialization on plastid function). The cytosolic GluRS was excluded from subsequent analyses because gain of mt targeting for this enzyme appears to be based on addition of a transit peptide by alternative splicing. Therefore, there is no change in the gene-body sequence relative to the enzyme that retains function in the cytosol. Final trimmed alignments and labeled trees are available as supplemental material (File S2).

Each aaRS sequence alignment and associated tree were used to perform branch-site tests for positive selection with three different tools in HyPhy v2.5.31 (Kosakovsky Pond, et al. 2020): aBSREL (Smith, et al. 2015), BUSTED (Murrell, et al. 2015), and MEME (Murrell, et al. 2012). Each analysis was performed twice – both with and without the “--branches Foreground” option. In addition to the MEME selection test, the output from this analysis was also used to perform a simple branch test because the MEME runs provide maximum likelihood values for when the entire tree is constrained to a single *d*_N_/*d*_S_ value and for when foreground and background are assigned two different *d*_N_/*d*_S_. These values were compared with a likelihood ratio test to determine whether allowing different *d*_N_/*d*_S_ ratios for the two tree partitions results in a significantly improved fit to the data.

## Data Availability

All datasets including expression TPM values, orthogroups sizes, and alignments and trees for rate analyses are available via https://github.com/dbsloan/aaRS_rates.

## ACKNOWLEDGEMENTS

This work was supported by a grant from the National Science Foundation (MCB-2048407).

